# ALK Inhibition Prolongs Survival in a Mouse Model of ALK-positive Anaplastic Thyroid Cancer

**DOI:** 10.1101/2025.03.25.642535

**Authors:** Yara M Machlah, Tim Brandenburg, G Sebastian Hönes, Sarah Theurer, Adrian D Prinz, Christoph Hoppe, Feyza Cansiz, Jens T Siveke, Johannes H. Schulte, Jukka Kero, Hendrik Undeutsch, Johannes Köster, Dagmar Führer, Lars C Moeller

## Abstract

**Background:** Anaplastic thyroid cancer (ATC) is the most aggressive thyroid cancer with a median survival of about 6 months. So far, no therapies offering a survival benefit are established. Thus, new therapeutic approaches are urgently needed. In general, genetic alterations leading to ATC increase PI3K and MAPK/ERK signalling and include mutations in receptor tyrosine kinases and tumour suppressor genes. They often occur together with the loss of P53, the most prevalent mutation in human ATC. Among such mutations are mutations and rearrangements of the *anaplastic lymphoma kinase* (ALK) gene.

**Methods:** To study ATC and potential treatment options, we generated a mouse model with inducible thyrocyte-specific expression of constitutively active mutant ALK^F1174L^ and homozygous deletion of *Trp53* due to a Cre recombinase under control of the thyroglobulin promoter (Tg-Cre^ERT2+/0^;LSL-ALK^F1174L/+^;Trp53^LoxP/LoxP^ mice, here referred to as Trp53^KO^/ALK^F1174L^ mice). Moreover, we established several primary thyroid cancer cell lines harbouring ALK^F1174L^ and Trp53^KO^ and investigated the effects of ALK inhibition *in vitro* and *in vivo*.

**Results:** Median survival of Trp53^KO^/ALK^F1174L^ mice was severely reduced and the mice showed massively enlarged thyroids. Histopathology confirmed development of locally invasive and metastatic ATC. Treatment of primary Trp53^KO^/ALK^F1174L^ ATC cells with the ALK inhibitor TAE-684 decreased AKT and ERK phosphorylation and induced a dose-dependent cytotoxicity. Trp53^KO^/ALK^F1174L^ mice treated with TAE-684 showed significantly extended median survival compared to the solvent group (66 days vs. 18 days, *p* < 0.0001).

**Conclusion:** Our data demonstrate that the combination of ALK^F1174L^ mutation with *Trp53* loss leads to the development of ATC. This study provides first functional data supporting the use of ALK inhibitors in patients with ALK-driven ATC. Our novel ATC mouse model and the derived cell lines offer valuable tools to explore the molecular characteristics of ATC, especially signalling pathway activation and tumour microenvironment, and to test novel therapeutics for the treatment of advanced thyroid cancers.

## Introduction

Anaplastic thyroid cancer (ATC) is among the most aggressive human malignancies with a median survival of less than 6 months (Lin et al., 2019). Although it accounts for less than 2% of thyroid carcinomas (TC), it is responsible for up to 50% of TC mortality (Alhejaily et al., 2023). ATC is characterised by a substantial mutational burden and genetic complexity. While oncogenic drivers like BRAF and RAS mutations are frequently detected in both differentiated and anaplastic TC, genetic defects affecting the tumour suppressor gene *Tp53*, the *TERT* promoter and members of the PI3K/AKT pathway are highly prevalent in ATC. Loss of function of *Tp53* constitutes the most frequent genetic defect in ATC and is involved in tumour dedifferentiation and progression. Over 70% of ATC cases harbour *Tp53* mutations, compared to only 8% in poorly differentiated thyroid cancer (PDTC) (Landa et al., 2016) and less than 1% in papillary thyroid cancer (PTC) (Cancer Genome Atlas Research, 2014).

Translocations involving the anaplastic lymphoma kinase (ALK) locus as well as activating ALK point mutations have been reported in various cancer types, including thyroid cancer, and result in an aberrant kinase activation (Godbert et al., 2015; Kelly et al., 2014; Murugan and Xing, 2011; Panebianco et al., 2019). ALK gene rearrangements, leading to fusions most frequently with striatin (STRN) or echinoderm microtubule-associated protein-like 4 (EML4), were detected in PTC (1-3%) as well as in the more aggressive TC types, PDTC (0-9%) and ATC (0-4%) (Chou et al., 2015; Kelly et al., 2014). The most frequent ALK mutations are the ligand-independent gain-of-function mutations F1174 and R1275, that reside within the ALK tyrosine kinase domain and comprise 85% of all ALK mutations (Holla et al., 2017). In ATC, Murugan and Xing identified two further gain-of-function ALK mutations, L1198F and G1201E, with a prevalence of 11.1% (2 of 18 ATCs). Both ALK mutants were shown to cause aberrant activation of the MAPK and PI3K/AKT pathways and promote cell transformation and invasion in *in vitro* assays (Murugan and Xing, 2011). Given that the dual activation of the two pathways driven by genetic alterations is a fundamental mechanism in the pathogenesis of ATC, activating ALK point mutations become relevant for thyroid tumorigenesis.

We have previously shown that overexpression of the activating ALK mutant F1174L in thyrocytes with subsequently increased MAPK and PI3K/AKT signalling result in the development of thyroid tumours resembling human PDTC (Kohler et al., 2019), which occupies an intermediate position between PTC and ATC regarding histopathological characteristics and clinical outcomes (Tong et al., 2022). In these ALK^F1174L^ mice, loss of follicular structure following tamoxifen injection induced overt hypothyroidism which, although not affecting the histological features of the tumours, promoted their progression and was associated with decreased median survival compared to ALK^F1174L^ mice kept euthyroid through L-thyroxine supplementation (Kohler et al., 2019). Nikitski et al. also demonstrated that thyroid-specific expression of the *STRN-ALK* fusion leads to tumours closely recapitulating PDTC (Nikitski et al., 2018), and that the combination of *STRN-ALK* fusion with biallelic *Trp53* loss results in progression to ATC (Nikitski et al., 2019). However, it remains unclear whether the combination of an ALK mutation with *p53* loss is sufficient to drive the development of ATC. If this combination indeed leads to ATC, the question arises as to whether these resulting tumours can be treated effectively with an ALK inhibitor.

In this study, we expanded our prior mouse model of thyroid-specific expression of the activating ALK mutation, F1174L, to investigate the impact of combining ALK^F1174L^ with a thyroid-specific inactivation of *Trp53* on thyroid tumorigenesis.

## Materials and Methods

### Generation of Trp53^KO^/ALK^F1174L^ mice and treatment

All animal studies were approved by the Landesamt für Natur, Umwelt und Verbraucherschutz Nordrhein-Westfalen (LANUV, Germany). Mice were housed under standard laboratory conditions (temperature 22 ±1°C and alternating 12-hour light and 12-hour dark cycle). Standard chow (Sniff, Soest, Germany) and tap water were provided *ad libitum*.

Mice expressing the lox-stop-lox (LSL)-ALK^F1174L^ mutant were previously established (Heukamp et al., 2012). Briefly, a genetic construct consisting of a synthetic CAG promoter, a transcriptional stop cassette flanked by loxP sites (lox-stop-lox model) and a constitutively active human ALK mutant, ALK^F1174L^, were introduced into the Rosa26 locus of C57BL/6 embryonic stem cells. An internal ribosomal entry site (IRES) and a construct encoding firefly luciferase are located downstream of the recombination site and enable the coupling of luciferase expression with ALK^F1174L^ expression.

To obtain mice with concomitant thyrocyte-specific heterozygous expression of ALK^F1174L^ and homozygous deletion of *Trp53,* triple mutants were generated. We crossed mice homozygous for the LSL-ALK^F1174L^ mutation and carrying exons 2-10 of *Trp53* flanked by loxP sites (LSL-ALK^F1174L^;Trp53^LoxP/LoxP^) with *Trp53*-floxed mice carrying tamoxifen-inducible Cre recombinase under the control of thyroglobulin (Tg) gene promoter (Tg-Cre^ERT2+/0^;Trp53^LoxP/LoxP^, here referred to as Trp53^KO^/ALK^WT^ mice). This resulted in Tg-Cre^ERT2+/0^;LSL-ALK^F1174L/+^;Trp53^LoxP/LoxP^ mice, here referred to as Trp53^KO^/ALK^F1174L^ mice. To activate the Cre recombinase, 7-week-old mice with a body weight > 20 g (male) or > 15 g (female) were administered a daily dose of 1 mg tamoxifen dissolved in 100 µl corn oil via i.p. injection for five consecutive days (5 mg in total) (Figure S1). From the time of tamoxifen injection all mice were supplied with drinking water containing L-thyroxine (LT4) at a concentration of 133 ng/mL (renewed two times per week) to maintain euthyroidism, avoid TSH effects on tumorigenesis and minimize interindividual differences in thyroid hormone levels. LT4 stock solution was prepared as previously described (Kohler et al., 2019). Mice were euthanized when predefined termination criteria, such as a loss of ≥ 20% bodyweight or significant difficulty in breathing, were met. The scoring system used to assess the condition of the mice is detailed in Table S1.

The ALK inhibitor, TAE-684 (Axon Medchem), was suspended in a solution consisting of 10% 1-methyl-2-pyrrolidinone and 90% PEG 300 (polyethylene glycol, molecular weight 300) (Sigma). Trp53^KO^/ALK^F1174L^ mice received either TAE-684 at a dose of 10 mg/kg or the solvent (10% 1-methyl-2-pyrrolidinone and 90% PEG 300) once daily via oral gavage for 30 consecutive days.

For subcutaneous tumour implantation, 1 x 10^6^ Trp53^KO^/ALK^F1174L^ mouse-derived thyroid cancer cells were resuspended in 50 µl culture medium and injected subcutaneously into the flank of C57BL/6 mice (Jackson Laboratory; 000664).

### PCR genotyping and confirmation of ALK recombination

Mice genotyping was performed using a polymerase chain reaction (PCR) program and the following pairs of primers for detection of LSL-ALK^F1174L^, TgCre^ERT2^, Trp53^LoxP^ and wild-type Trp53: ALKF1174L-KI-rev 5’-CCC AAG GCA CAC AAA AAA CC-3’, ALKF1174L-KI-fwd 5’-TGG CAG GCT TGA GAT CTG G-3’, Cre-ERT2-rev 5’-TGG CAG CTC TCA TGT CTC CAG-3’, Cre-ERT2-fwd 5’-TCA GAG ATA CCT GGC CTG GTC-3’, Trp53flox-rev 5’-GGA GGC AGA GAC AGT TGG AG-3’, Trp53flox-fwd 5’-GGT TAA ACC CAG CTT GAC CA-3’. Successful Cre-mediated ALK recombination was confirmed by PCR with the following primers: ALK-recom-rev: 5’-GCA CCA CGA AGT CAA CTG C-3’, ALKF1174L-fwd: 5’-TGG CAG GCT TGA GAT CTG G-3’, Rosa26-wt-rev: 5’-CAT GTC TTT AAT CTA CCT CGA TGG-3’, Rosa26-wt-fwd: 5’-CTC TTC CCT CGT GAT CTG CAA CTC C -3’.

### Cell line establishment and culture

To establish primary thyroid cancer cell lines from Trp53^KO^/ALK^F1174L^ mice, half of the thyroid tumour was dissected for culturing and stored on ice. The remaining tumour tissues were fixed in 4% PFA or snap frozen for further analyses. The tumour material was rinsed using a wash solution consisting of advanced DMEM/F-12 (Gibco™, Thermo Fisher Scientific, Darmstadt) supplemented with 1% GlutaMAX (Gibco™, Thermo Fisher Scientific), 1% HEPES (Gibco™, Thermo Fisher Scientific), and Primocin at 125 µl/50 ml (InvivoGen). The tumour tissue was then cut into small fragments, which were mixed for 15 minutes at 37°C in a digestion solution containing Collagenase Type II (50 mg/1 ml; Thermo Fisher Scientific) and Dispase II (12.5 mg/1 ml; Thermo Fisher Scientific). The digestion step was repeated as needed until the remaining tissue pieces were completely disaggregated. The supernatant containing dissociated cells was then transferred to a wash solution and centrifuged at 130 × g at 4°C. Primary cell pellets were subsequently suspended in a culture medium composed of advanced DMEM/F-12 containing 1% GlutaMAX, 1% HEPES, and 10% fetal bovine serum (Gibco™, Thermo Fisher Scientific). Immediately before use, 125 µl of primocin were added to every 50 ml of culture medium. Cell cultures were maintained under standard conditions in a humidified atmosphere (5% CO_2_, 37°C) and harvested using 0.25% trypsin/EDTA for cell passaging. Cell counting was performed using trypan blue (Sigma Aldrich) and a hemocytometer. For cell culture experiments, TAE-684 (Axon Medchem) was dissolved in DMSO at a concentration of 5 mM.

### Bioluminescence imaging

Luciferase activity in Trp53^KO^/ALK^F1174L^ mouse-derived thyroid cancer cells was detected using an IVIS Lumina II imaging system after treatment with 150 µg/ml of D-luciferin (Perkin Elmer) dissolved in PBS for 5 minutes.

### Sulforhodamine B (SRB) assay

To measure drug-induced cytotoxicity we used the Sulforhodamine B (SRB) assay (Skehan et al., 1990). Primary thyroid cancer cells generated from the Trp53^KO^/ALK^F1174L^ mouse model were seeded in 96-well microplates at densities of 5,000 cells in 200 µl of culture medium per well. 24 hours after seeding, cells were treated with DMSO (negative control) or increasing concentrations of TAE-684 for 48 hours. Cultures were then fixed with trichloroacetic (TCA) for 1 hour at 4°C at a final TCA concentration of 10%. Subsequently plates were washed four times with tap water, air dried and stained for 30 minutes with 0.4% SRB dissolved in 1% acetic acid. Unbound dye was removed by four washes with 1% acetic acid. After being air dried, protein-bound dye was solubilized with 10 mM Tris base (pH 10.5) for 10 minutes on a gyratory shaker. Absorbance was measured at 565 nm with a microplate reader. Background optical densities were measured in wells with culture medium without cells. Dose-response curves, expressed as mean values with standard error, and EC_50_ values were determined with GraphPad Prism (version 10.0.3 (217) for macOS).

### Histopathological examination and immunochemistry

Histological analysis was performed on formalin-fixed, paraffin-embedded (FFPE) blocks. Tissue blocks were cut at thickness of 4 µm, deparaffinized, and stained with hematoxylin and eosin (H&E) or immunohistochemically (IHC). For IHC, sections were boiled in Tris/EDTA buffer, pH 9 (Dako, S2367) for 20 minutes to accomplish antigen retrieval. Slides were incubated for 30 minutes with primary antibodies against: thyroglobulin (1:100; abcam; ab156008), Nkx2.1 (1:100; MA5-13961; Invitrogen), PAX8 (1:500; abcam; ab239363) and TPO (1:1000; abcam; ab278525). Immunodetection was performed with the anti-rabbit Polyview Plus AP (Enzo Life Sciences; ENZ-ACC110-0150) and Permanent AP Red (Zytomed; ZUC001-125). Nuclei were stained with hematoxylin (Zytomed; ZUC087-500).

Mouse thyroid and lung sections were examined by an experienced thyroid pathologist blinded to the genotype. Diagnoses of PTC and ATC were made according to the current criteria, published at WHO classification for endocrine tumours (Baloch et al., 2022).

### Immunoblotting

Whole-protein lysates were obtained from primary thyroid cancer cells or from snap-frozen thyroid tissue using RIPA buffer (150 mM NaCl, 50 mM HCl, 1% Nonidet P-40, 0.5% sodium deoxycholate, 0.1% SDS, 2 mM EDTA, 50 mM NaF) supplemented with PhosSTOP and cOmplete Protease Inhibitor Cocktail (Sigma-Aldrich). After 20 minutes of centrifugation with 17,000 × g at 4°C, the supernatant was collected, and the cell debris was discarded. A total of 20 µg of protein were separated by SDS/PAGE and transferred onto a PVDF membrane (Roti-Fluoro PVDF; Roth) overnight at 4°C. After blocking with 5% BSA in TBS-T (Tris-buffered saline with 0.5% Tween-20) for 1 h membranes were incubated for 16 h at 4°C under gentle agitation with the desired primary antibody against *phospho-p44/42 MAPK* (1:1000, #4370; Cell Signaling), *p44/42 MAPK* (Erk1/2) (1:1000, #4695; Cell Signaling), *phospho-AKT* (Thr308) (1:1000, #2965; Cell Signaling), and *AKT* (1:1000, #9272; Cell Signaling). HRP-conjugated secondary antibody against rabbit IgG (1:2000; #7074 Cell Signaling) served as a second antibody. The signals were detected with a VersaDocMP4000 (BioRad) and the band densities were determined with Image Lab software (BioRad).

### Gene expression analysis and RNA sequencing

Total RNA was isolated from the thyroid gland or primary thyroid cancer cells using the RNeasy Kit (Qiagen) and stored at -80°C. Subsequently, 2 µg of total RNA were reverse transcribed into cDNA using SuperScript III (Invitrogen) and random hexamer primers. Quantitative real-time PCR (qRTPCR) was performed using Roche SYBR Green I Master Mix on a LightCycler LC480 (Roche). Primer sequences will be provided at request. The expression was normalised to that of 18S RNA. The fold change in gene expression was calculated using the efficiency-corrected method (Pfaffl, 2001).

For total transcriptome analysis, concentration and quality of RNA were measured with Qubit (Invitrogen, Waltham, MA, USA) and Agilent Bioanalyzer (Agilent, SantaClara, CA, USA). Library preparation was performed using Lexogens QuantSeq 3’ mRNA-Seq Library Prep Kit FWD (Lexogen, Inc., Greenland, NH, USA) and quantified with Library Quant qPCR. Samples were sequenced on a NextSeq500 (Illumina, San Diego, CA, USA). Sequences were trimmed with TrimGalore (Martin, 2011) and aligned with hisat2 (Kim et al., 2019) to hg38. After adapter removal with Cutadapt (Martin, 2011), transcripts were quantified with Kallisto (Bray et al., 2016). Integrated normalization and differential expression analysis was conducted with Sleuth (Pimentel et al., 2017).

### Statistical methods

All statistical analyses were performed using GraphPad Prism version 10.0.3 (217) for macOS. Difference in survival was assessed by a log-rank test. If not specified, parametric data were compared with a two-tailed *t* test and presented as the mean ± SD. *P* values of < 0.05 were considered statistically significant.

## Results

### Trp53^KO^/ALK^F1174L^ mice exhibit reduced survival and massively enlarged thyroid glands

Thyroid-specific heterozygous expression of ALK^F1174L^ and biallelic *Trp53* loss led to severe decrease in the survival of Trp53^KO^/ALK^F1174L^ mice (*n=31*), with a median survival of 105 days, compared to control Trp53^KO^/ALK^WT^ mice (*n=17*), which survived the observation period without complications (*p* < 0.0001) (Figure 1A). Inspection and manual palpation of the necks of Trp53^KO^/ALK^F1174L^ mice revealed large hard cervical masses firmly attached to the surrounding tissues. A massive increase in volume and a tumorous transformation of the thyroid gland were confirmed after dissection (Figure 1B). Compared to Trp53^KO^/ALK^WT^ mice, Trp53^KO^/ALK^F1174L^ mice showed an up to 100-fold higher thyroid weight (average gland weight 288.5 ±206.9 mg, vs. 13.5 ±7.9 mg in controls; *p* < 0.0001). We observed a sex-specific difference in survival, with a shorter median survival of 75 days in male Trp53^KO^/ALK^F1174L^ mice (*n=14*), while females survived for a median of 116 days (*n=17*) (*p* = 0.0023) (Figure 1C). No difference in thyroid weight was observed between male and female Trp53^KO^/ALK^F1174L^ mice (Figure 1D).

**Figure 1.**
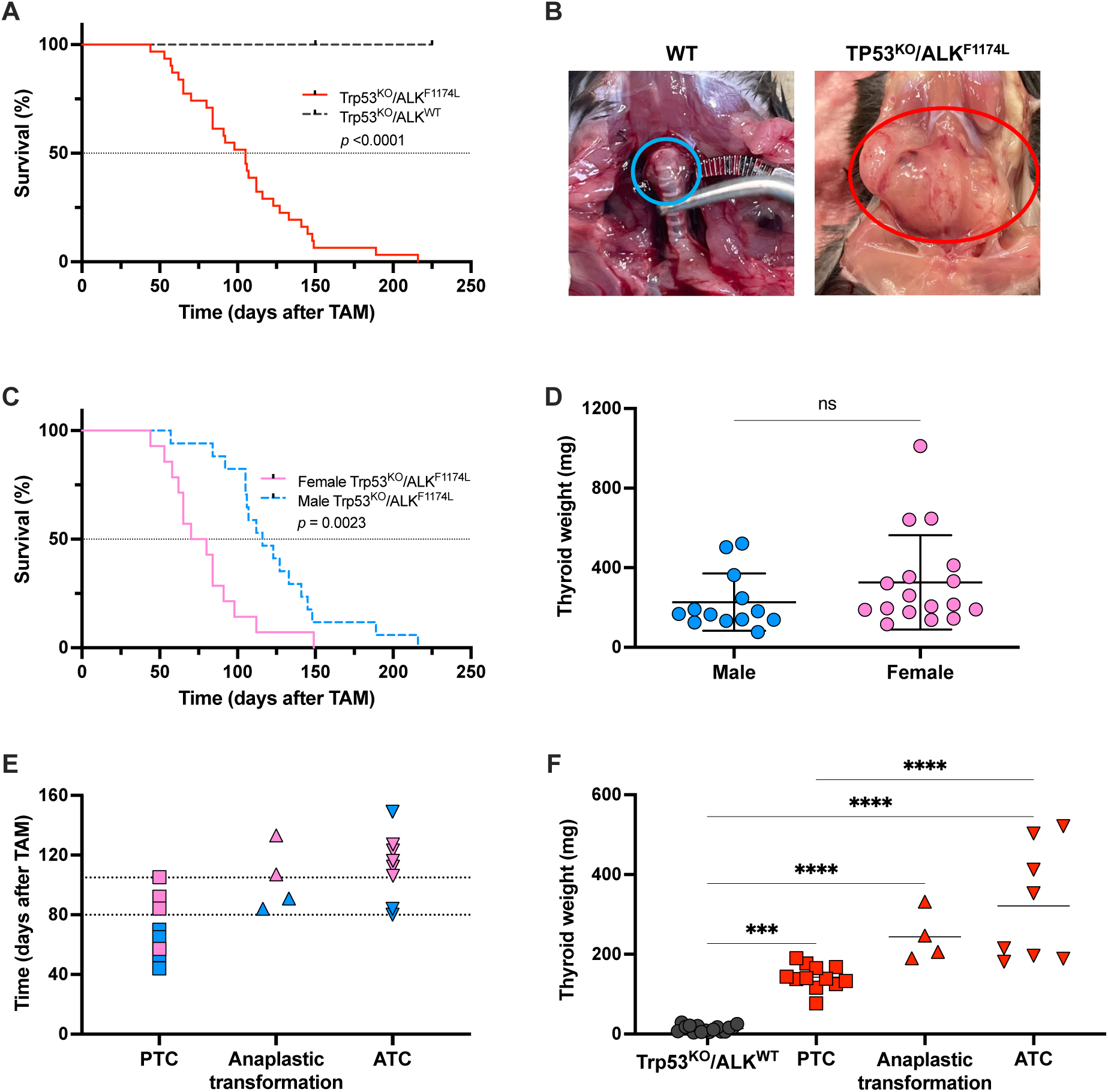
Survival and thyroid tumour development in Trp53^KO^/ALK^F1174L^ mice. (**A**) Kaplan-Meier survival curve of Trp53^KO^/ALK^F1174L^ mice (*n=31*) compared to Trp53^KO^/ALK^WT^ mice (*n=17*) after tamoxifen (TAM) administration. (**B**) Representative images of thyroid tumours in Trp53^KO^/ALK^F1174L^ mice (bottom row, circled in red) compared to normal thyroid (top row, circled in blue); WT, wild type. (**C**) Survival of male Trp53^KO^/ALK^F1174L^ mice (*n=17*) vs. female Trp53^KO^/ALK^F1174L^ mice (*n=14*). (**D**) Comparison of thyroid weights between male and female Trp53^KO^/ALK^F1174L^ mice; ns, not significant. (**E**) Number of days after tamoxifen (TAM) administration when papillary thyroid carcinoma (PTC), anaplastic transformation, and anaplastic thyroid cancer (ATC) were detected in male (blue) and female (pink) Trp53^KO^/ALK^F1174L^ mice. (**F**) Thyroid weight measured on the day of tissue collection; one-way ANOVA and Tukey’s test. ***, *p* < 0.001; ****, *p* < 0.0001.

### Thyroid-specific Trp53^KO^/ALK^F1174L^ leads to development of ATC and pulmonary metastases

All Trp53^KO^/ALK^F1174L^ mice developed histologically confirmed thyroid cancers: 12 PTCs, four PTCs with transition into ATC and eight ATCs. Well-differentiated PTCs were predominantly observed in the early stages following Cre activation, whereas the earliest ATCs were identified in male mice sacrificed on day 80 and female mice on day 105 after tamoxifen administration (Figure 1E). Regarding tumour weight, mice with ATC exhibited a significantly higher tumour weight (321.3 ±144.6 mg) than those that developed PTC (142.7 ±30.20 mg) (*p* = 0.0005) (Figure 1F). Histologically, PTCs exhibited microscopic features consistent with the classic and/or follicular subtype of papillary carcinoma (Figure 2A). In four cases, we noted the coexistence of PTC and ATC within the same thyroid gland, with the anaplastic component manifesting as small, distinct foci adjacent to the dominant PTC (Figure 2B). In ATC tumours, a spindle cell pattern was observed (Figure 2C, 2D). Immunohistochemical analysis of ATCs revealed loss of thyroid differentiation markers Tg, TTF-1 and TPO (Figure 2E - 2G). Trp53^KO^/ALK^F1174L^-driven tumours displayed an aggressive tumour behaviour leading to local invasiveness and distant metastasis. They invaded locally into the surrounding strap muscles and trachea (Figure 2C, 2D) and metastasized to the lungs (Figure 2H - 2J). Interestingly, we observed a discrepancy between the histological pattern of the thyroid tumour and the lung metastases within the same animal. Notably, PTC lung metastases were observed in mice with histologically confirmed ATC in the thyroid. Thyroid-specific homozygous Trp53 inactivation in Trp53^KO^/ALK^WT^ mice did not cause any overt phenotype and survival was not different from WT mice. Gene Ontology (GO) term enrichment analysis showed that differentially regulated genes in Trp53^KO^/ALK^F1174L^-driven thyroid tumours were primarily associated with cellular compartments such as the nucleus, membrane, and cytosol, as well as molecular functions including protein binding and metal ion binding (Figure 3A). Several genes associated with cancer progression, including RSPO4, CHST8, HECW1 and GAP43, were markedly elevated in Trp53^KO^/ALK^F1174L^-ATC compared to Trp53^KO^/ALK^F1174L^ -PTC. Additionally, genes related to epithelial-mesenchymal transition (SOX2) and genetic instability (ASXL3) showed significant upregulation in Trp53^KO^/ALK^F1174L^-ATC (Figure 3B).

**Figure 2.**
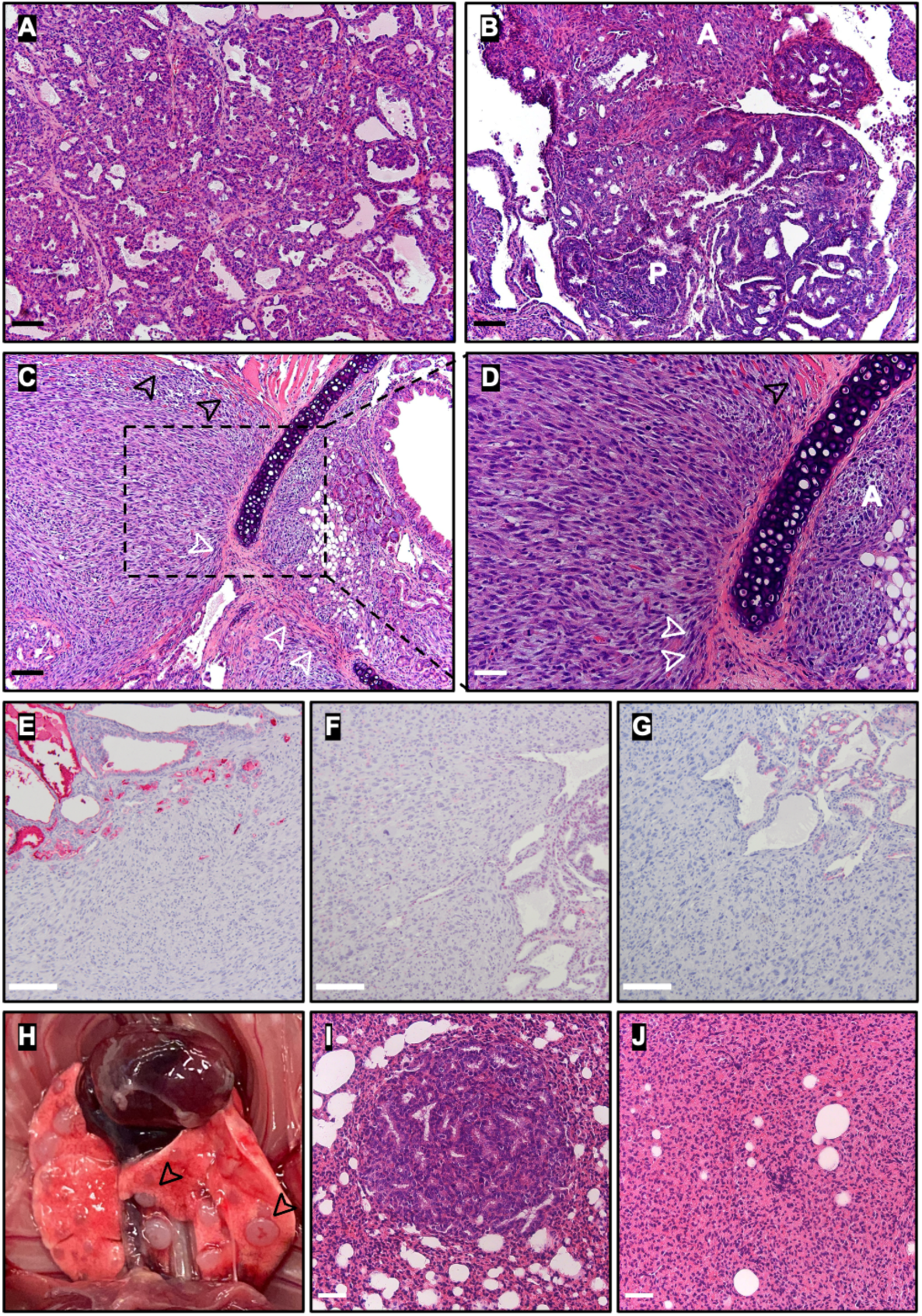
Locally invasive and metastatic PTC and ATC in Trp53^KO^/ALK^F1174L^ mice. (**A**) H&E staining of the thyroid of Trp53^KO^/ALK^F1174L^ mouse sacrificed on day 105 after tamoxifen administration, showing PTC. (**B**) Area of anaplastic transformation adjacent to PTC (H&E; A, ATC; P, PTC). (**C, D**) ATC in Trp53^KO^/ALK^F1174L^ mouse sacrificed on day 136, showing infiltration of surrounding strap muscles (black arrows) and tumour extension into submucosal areas through intercartilaginous portion (white arrows) (H&E; A, ATC). IHC showing loss of Tg (**E**), TTF1 (**F**) und TPO (**G**) in ATC compared to differentiated areas of the thyroid. (**H**) Photograph of lung tissue with metastatic lesions (black arrows). Lung metastases of PTC (**I**) and ATC (**J**) (H&E). Black scale bar = 100 µm; white scale bar = 50 µm.

**Figure 3.**
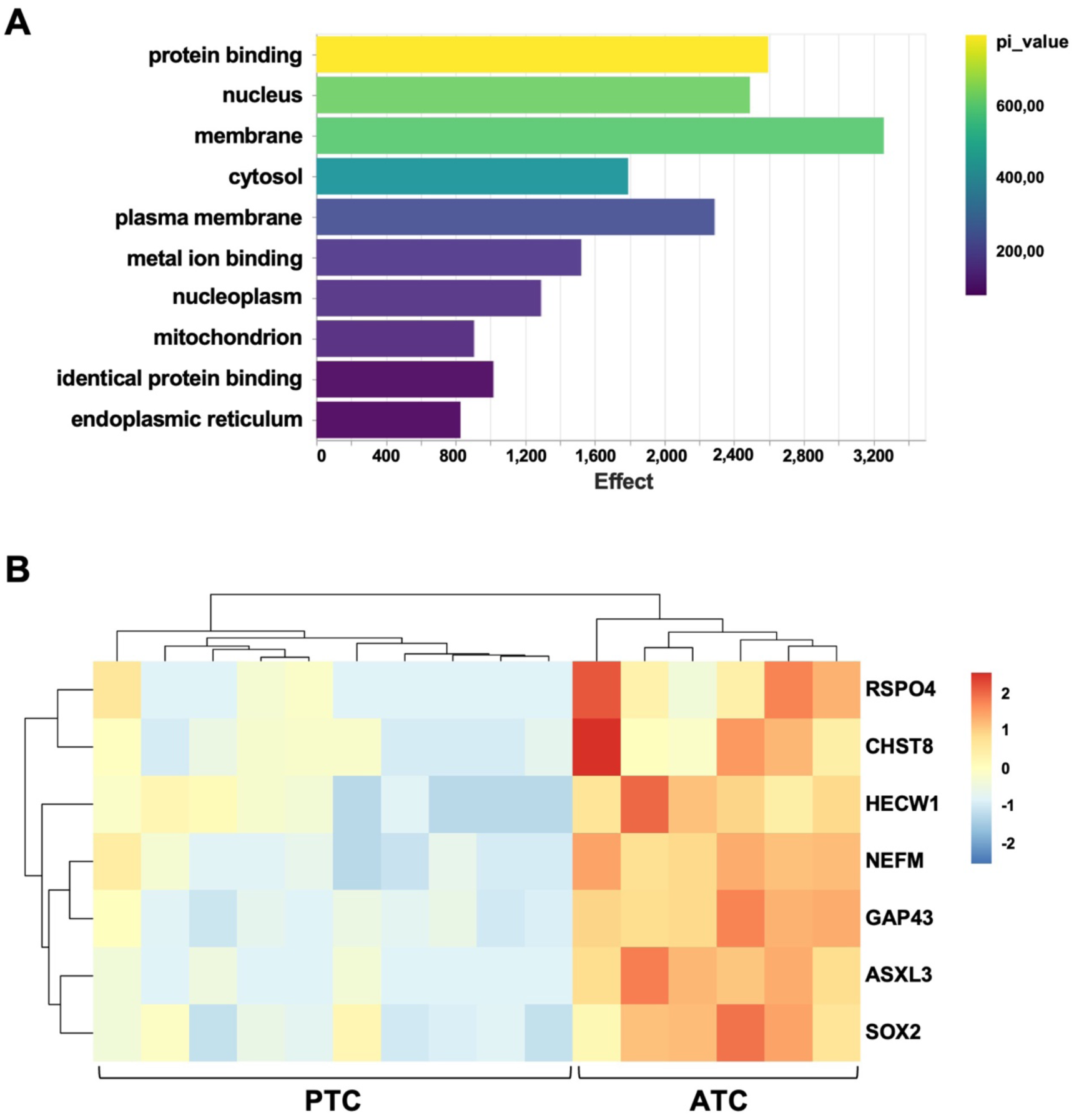
Gene ontology term enrichment and heatmap analysis of differentially expressed genes in Trp53^KO^/ALK^F1174L^ thyroid tumours. (**A**) Top 10 enriched GO terms based on differentially expressed genes in Trp53^KO^/ALK^F1174L^-driven thyroid tumours. (**B**) Heatmap of the most upregulated genes in Trp53^KO^/ALK^F1174L^ -ATC compared to PTC.

### Transplanted primary ALK-positive ATC cells grow to ATCs *in vivo*

To establish an *in vitro* model, we generated ten primary cell lines from thyroid tumours of Trp53^KO^/ALK^F1174L^ mice at various stages after Cre activation. Histological analysis of the corresponding in situ tumours revealed that six cell lines originated from PTC and four from ATC. To confirm that the primary cells derived from thyrocytes expressing the ALK^F1174L^ mutation, DNA was extracted from a cell pellet, and a PCR was conducted. The PCR results confirmed the Cre-mediated recombination in the cells (Figure 4A). Additionally, given that the ALK^F1174L^ transgene also encodes for luciferase, we treated the cell culture with luciferin and detected the bioluminescence signal (Figure 4B).

**Figure 4.**
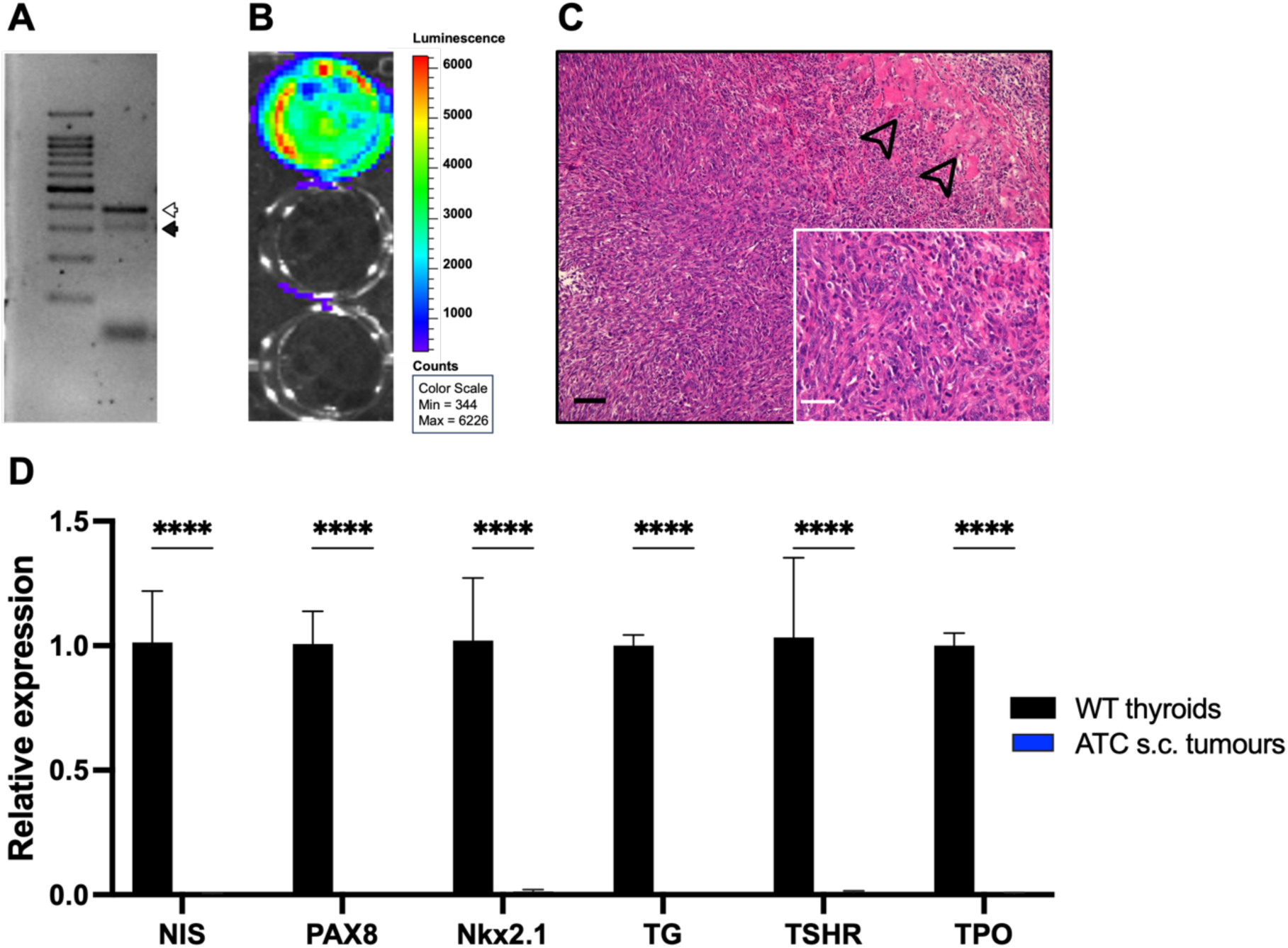
Primary thyroid cancer cells of Trp53^KO^/ALK^F1174L^ mice. (**A**) Confirmation of ALK recombination (open arrowhead) and Rosa26 locus (filled arrowhead) in cells derived from a Trp53^KO^/ALK^F1174L^ mouse through PCR on genomic DNA. (**B**) Measurement of bioluminescence signal from luciferase as a surrogate for ALK^F1147L^ expression. (**C**) H&E staining of the subcutaneous tumour induced by Tiara-D177 cells, showing ATC with invasive growth into surrounding muscle tissue (black arrows; black scale bar = 100 µm; white scale bar = 50 µm). (**D**) Gene expression analysis of s.c. ATC demonstrating loss of thyroid differentiation markers compared to normal thyroid tissue. ****, *p* < 0.0001.

To determine whether the newly established primary cells maintain histological features and growth patterns of the original thyroid tumours, one of the primary thyroid cancer cell lines (Tiara-D177), was subcutaneously (s.c.) injected into the flank of C57BL/6 mice (*n=3*). Tiara-D177 cells were derived from a female Trp53^KO^/ALK^F1174L^ ATC-bearing mouse sacrificed on day 177 after tamoxifen injection with a final thyroid weight of 1.086 mg. Following s.c. tumour implantation, the mice were regularly monitored for the occurrence of termination criteria. After three weeks, the s.c. tumours reached a diameter of 15 mm and the mice were sacrificed. The isolated s.c. tumours, induced by the injection of Tiara-D177 cells, were histologically identified as ATC and thus corresponded to the *in situ* thyroid tumour from which the Tiara-D177 cell line originated (Figure 4C). Gene expression analysis on Tiara-D177 s.c. tumours again demonstrated a complete loss of thyroid differentiation markers compared to normal thyroid tissue (Figure 4D).

### ALK inhibition induces cytotoxicity in Trp53^KO^/ALK^F1174L^ ATC cells *in vitro*

To investigate the effect of ALK inhibition in Trp53^KO^/ALK^F1174L^ ATC-cells, Tiara-D177 culture was evaluated for cell viability and cell signalling in response to the ALK inhibitor TAE-684. Tiara-D177 cells were incubated with serial dilutions of TAE-684 (15.6 nM to 4.0 µM) or solvent for 48 hours to measure drug-induced cytotoxicity. TAE-684 inhibited the proliferation of Tiara-D177 cells in a dose-dependent manner with an EC_50_ of 442.8 nM (Figure 5A). Furthermore, *in vitro* treatment with TAE-684 suppressed the ALK-induced phosphorylation of ERK and AKT compared to solvent treatment (Figure 5B - 5D). Similar results were observed in two further PTC cell lines, generated on day 74 and 105 after tamoxifen administration (Figure S2).

**Figure 5.**
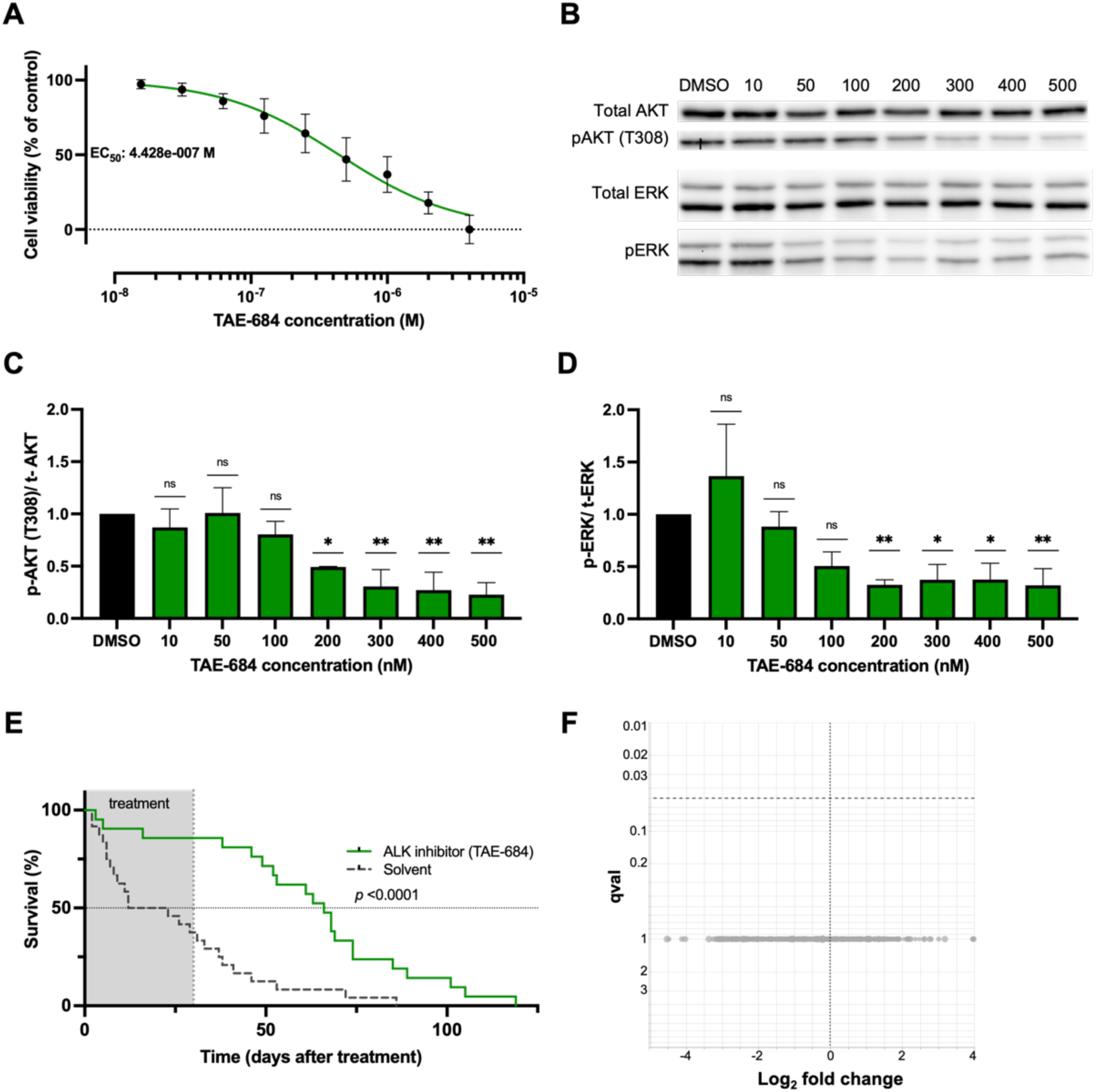
ALK inhibition in Trp53^KO^/ALK^F1174L^ cells and mice. (**A**) SRB assay of the Trp53^KO^/ALK^F1174L^-derived ATC cell line Tiara-D177 treated with TAE-684 assessing cell survival. Values normalised (100%) to vehicle-treated control (three independent experiments with eight biological replicates). (**B-D**) Western blot and densitometry results of phospho-AKT (T308), total AKT, phospho-ERK1/2 and total ERK1/2 in lysates from ATC Tiara-D177 cells treated with DMSO or TAE-684 in nM (three independent experiments with two biological replicates). (**E**) Kaplan-Meier survival curve showing prolonged survival in TAE-684-treated Trp53^KO^/ALK^F1174L^ mice (*n=21*) compared to solvent-treated mice (*n=24*). (**F**) Volcano plot for the comparison between the TAE-684-treated and solvent-treated thyroid tumours. * = *p* < 0.05; ** = *p* < 0.01; ns, not significant.

### ALK inhibition prolonged survival *in vivo*

After analysing the first cohort of Trp53^KO^/ALK^F1174L^ mice for survival and histological characteristics, a new mouse cohort was generated to examine the effects of ALK inhibition in Trp53^KO^/ALK^F1174L^-driven thyroid cancer (*n=45*). 21 mice received TAE-684 and 24 mice were administered the vehicle solution for 30 consecutive days. The start of the 30-day treatment period was determined based on the time point of the detection of the earliest ATC in the previously analysed survival cohort. Thus, treatment was initiated 80 days after tamoxifen injection male and 105 days after tamoxifen injection in female mice. Throughout the experiment, mice were closely monitored and removed from the experiment upon reaching predefined termination criteria. The corresponding survival curve showed a significant survival advantage for the inhibitor-treated group compared to the solvent-treated control group (66 days vs. 18 days, *p* < 0.0001) (Figure 5E). Over the course of the treatment period, body weight stabilisation was observed in mice subjected to TAE-684 (Figure S3 A-B). With the exception of three mice (two of which were found dead in their cages), TAE-684-treated mice survived the full 30-day treatment. In contrast, over half of the control mice (*n=15*) met termination criteria within 30 days. Final thyroid gland weight did not vary between vehicle- and TAE-684-treated groups (Figure S3 C). Retrospective histological examination revealed that approximately 50% of both the TAE-684-treated mice (*n=10*) and the control cohort (*n=11*) had ATC. RNA sequencing data showed no significant transcriptome differences between solvent- and TAE-684-treated tumours, which were collected after the end of ALK inhibition (Figure 5F). The occurrence of lung metastases was similar in mice treated with TAE-684 (5 of 19 mice) compared to those treated with solvent (3 of 24 mice) (*p* = 0.432 in Fisher’s exact test).

## Discussion

ATC is a rare yet highly lethal malignancy. The dismal prognosis of ATC results from its rapid progression and limited response to current therapy modalities. The genetic complexity and heterogeneity that characterise ATC impede identifying new therapeutic targets and thus developing effective treatments. Despite scientific and clinical advances in this field including tyrosine kinase inhibitors and immunotherapy there remains an urgent need for more effective research and therapeutic approaches (Bible et al., 2021). Most functional data regarding the molecular background of ATC predominantly corroborate the role of BRAF^V600E^ mutation as an initiator in thyroid carcinogenesis (Landa and Knauf, 2019). The involvement of less prevalent genetic alterations in the onset of thyroid carcinomas has not been thoroughly investigated thus far.

In prior work, we demonstrated that the thyrocyte-specific overexpression of the activating ALK^F1174L^ mutation, that leads to the activation of MAPK and PI3K signalling pathways, results in the development of PDTC. However, it is not sufficient to induce complete tumour dedifferentiation (Kohler et al., 2019). Genomic studies of human TC specimens emphasise the role of Tp53 as a tumour suppressor in TC progression. Mutations in *Tp53* are frequently identified in human ATC, less frequent in PDTC, and rarely in PTC (Cancer Genome Atlas Research, 2014; Landa et al., 2016).

Building upon these data, we generated a novel spatiotemporally controlled mouse model of ATC, obtained by combining two molecular alterations: activation of ALK (ALK^F1174L^ mutation) and inactivation of *Trp53* (deletion of exons 2-10). ALK^F1174L^/Trp53^KO^-driven tumours closely recapitulate features of advanced human thyroid cancer with evidence of extrathyroidal extension, lung metastases and a lethal outcome.

By comparing both mouse models, ALK^F1174L^ and Trp53^KO^/ALK^F1174L^ mice, it becomes evident that an activating ALK mutation can initiate thyroid tumorigenesis on its own (ALK^F1174L^ -> PDTC) but is insufficient for anaplastic transformation. On the other hand, thyroid-specific *Trp53* loss, while incapable of driving tumorigenesis independently (Trp53^KO^/ALK^WT^ -> no phenotype changes), acts as a prerequisite for thyroid cancer progression into ATC, and only results in ATC development when combined with another driver mutation such as the ALK^F1174L^ mutation (Trp53^KO^/ALK^F1174L^ -> ATC). This aligns with other mouse models that have demonstrated that inactivation of *Tp53* promotes tumour progression from well-differentiated TC into ATC when combined with a *BRAF* mutation (McFadden et al., 2014), *Pten* deletion (Antico Arciuch et al., 2011) or *RET/PTC* fusion (La Perle et al., 2000).

Our data support a stepwise thyroid carcinogenesis model with progression from well-differentiated to undifferentiated TC. First, in the early phase after tamoxifen administration, we predominantly detected PTCs and the frequency of ATC increased over time. Second, histological analyses revealed areas of anaplastic transformation alongside PTC within the same thyroid gland. Third, RNA sequencing data showed upregulation of several genes in ATC compared to PTC, which are linked to tumour progression (Chou et al., 2024; da Silva et al., 2022; Liu et al., 2022; Shukla et al., 2017; Zhang et al., 2018), dedifferentiation (Herreros-Villanueva et al., 2013) and epithelial-mesenchymal transition (EMT) (Han et al., 2012) (Gao et al., 2015). In addition, some mice with ATC had lung metastases with the histology of PTC, suggesting that these ATCs originated from pre-existing PTCs. This aligns with next-generation sequencing data of human ATC, which suggests that ATCs evolve from their well-differentiated counterparts (Landa et al., 2016). Although the expression of ALK^F1174L^ and *Trp53* inactivation through tamoxifen was simultaneously induced from the outset, a certain temporal latency was required until the complete ATC phenotype was achieved. Whether the dedifferentiation in ALK^F1174L^/Trp53^KO^-driven tumours occurs by acquiring additional genetic or epigenetic alterations requires more functional in-depth data.

Due to the experimental design with survival as the primary endpoint, the precise timing of the onset of anaplastic transformation in Trp53^KO^/ALK^F1174L^ mice cannot be determined accurately, as the mice were sacrificed based on predefined criteria rather than at specified time intervals. However, histological analysis confirmed the presence of a complete ATC phenotype as early as 80 days after Cre-mediated tumour induction. Consistent with this, Trp53^KO^/ALK^F1174L^ mice exhibit an extremely shortened median survival of 105 days post tamoxifen. In comparison, a median survival of approximately 6 months after the induction of thyrocyte-specific BRAF^V600E^ expression and *p53* loss was observed in a BRAF^V600E^-driven ATC mouse model (McFadden et al., 2014). Another *in vivo* ATC model, driven by the combined loss of *Pten* and *p53*, shows a longer ATC latency, despite using an embryonic onset of the Cre-recombinase. Mice with homozygous loss of *Pten* and *p53* had a median survival of 38.4 weeks, and over 75% of them developed ATC by 9 months of age. However, after 4 to 8 months only well-differentiated carcinomas were induced (Antico Arciuch et al., 2011). Taken together, Trp53^KO^/ALK^F1174L^ mice exhibit an acceleration of TC progression when compared to models with different genetic alterations. Due to variations in the experimental design, a direct comparison between mouse models cannot be drawn. However, these data imply that thyroid tumours induced by simultaneous ALK^F1174L^ mutation and Trp53^KO^ display more aggressive tumour behaviour. This may be attributed to the fact that constitutive ALK activation leads to aberrant activation of its downstream pathways including both MAPK and PI3K cascades (Jiang and Ji, 2019). Further data are required to verify whether the nature of the oncogenic event distinctly influences the biological behaviour of the resulting ATC. For this purpose, our mouse model provides a suitable tool.

Interestingly, we noted an extended median survival among female Trp53^KO^/ALK^F1174L^ mice compared to their male counterparts. In the female cohort, ATC was identified later than in the male cohort, and no sex-specific difference was observed in final tumour weight. This implies that the prolonged survival of female mice is not a genuine survival advantage but rather a temporal shift in the onset of transformation. A comparable finding was observed in a BRAF^V600E^ PTC mouse model by Knauf et al., with a higher frequency of dedifferentiation and vascular invasion in males (Knauf et al., 2005). Considering the distinct distribution of oestrogen receptors between males and females, the observed latency in Trp53^KO^/ALK^F1174L^ female mice could be associated with the tamoxifen-inducible Cre recombinase technology employed in this study. Sex differences in terms of metabolism and thermogenesis of adipose tissue upon tamoxifen administration have been reported (Zhao et al., 2020). To our knowledge, this phenomenon has not been documented in tamoxifen induced cancer mouse models.

Currently, limited information is available on the efficacy of ALK inhibitors in ATC (Godbert et al., 2015; Leroy et al., 2020). Previous studies have reported resistance of ALK^F1174L^-mutated cells to crizotinib. Therefore, in this study we used the ALK inhibitor TAE-684 (Berry et al., 2012; Heukamp et al., 2012). Our findings demonstrate that TAE-684 induces dose-dependent cytotoxicity in the ATC primary cell line harbouring ALK^F1174L^ and Trp53^KO^ and leads to decreased phosphorylation levels of downstream mediators of the ALK signalling pathway, AKT and ERK1/2. In addition, a thirty-day oral treatment of Trp53^KO^/ALK^F1174L^ transgenic mice with the ALK inhibitor TAE-684 resulted in a significant survival benefit compared to the solvent-treated cohort.

The fact that most of the inhibitor-treated animals survived throughout the treatment period and only deteriorated after therapy discontinuation indicates a drug-dependent effect, which likely led to regression or stabilisation of the tumours during therapy. This is further supported by the observation that transcriptome data and thyroid tumour weights collected after treatment completion did not differ between the inhibitor- and solvent-administered cohorts, despite the prolonged survival of the inhibitor-treated mice. The susceptibility of cells and tumours carrying ALK^F1174L^ and Trp53^KO^ to TAE-684 implicates their oncogene addiction to ALK^F1174L^. Notably, ALK seems to persist as a driving oncogene, irrespective of the presence of additional genetic alterations.

In conclusion, we present a novel mouse model of ATC and provide evidence that the combination of ALK^F1174L^ mutation with *Trp53* loss leads to the development of ATC, which recapitulates features of aggressive human thyroid cancer. This study offers first functional data supporting the use of ALK inhibitors in patients with ALK-driven ATC. We propose this model as a suitable experimental tool for further understanding of molecular events associated with thyroid tumour dedifferentiation and progression. Moreover, this model can be used for testing novel diagnostic and therapeutic strategies of ATC.

## Acknowledgments

We thank Andrea Jaeger and Julius Göbel (Department of Endocrinology, Diabetes and Metabolism, University Hospital Essen) for their excellent technical assistance. Moreover, we thank the Imaging Center Essen, a Service Core Facility of the Faculty of Medicine of the University Duisburg-Essen, for providing Caliper IVIS Lumina II and the Genomics and Transcriptomics Facility (University Hospital Essen) for performance of RNA-sequencing.

## Supplementary

**Figure S1.**
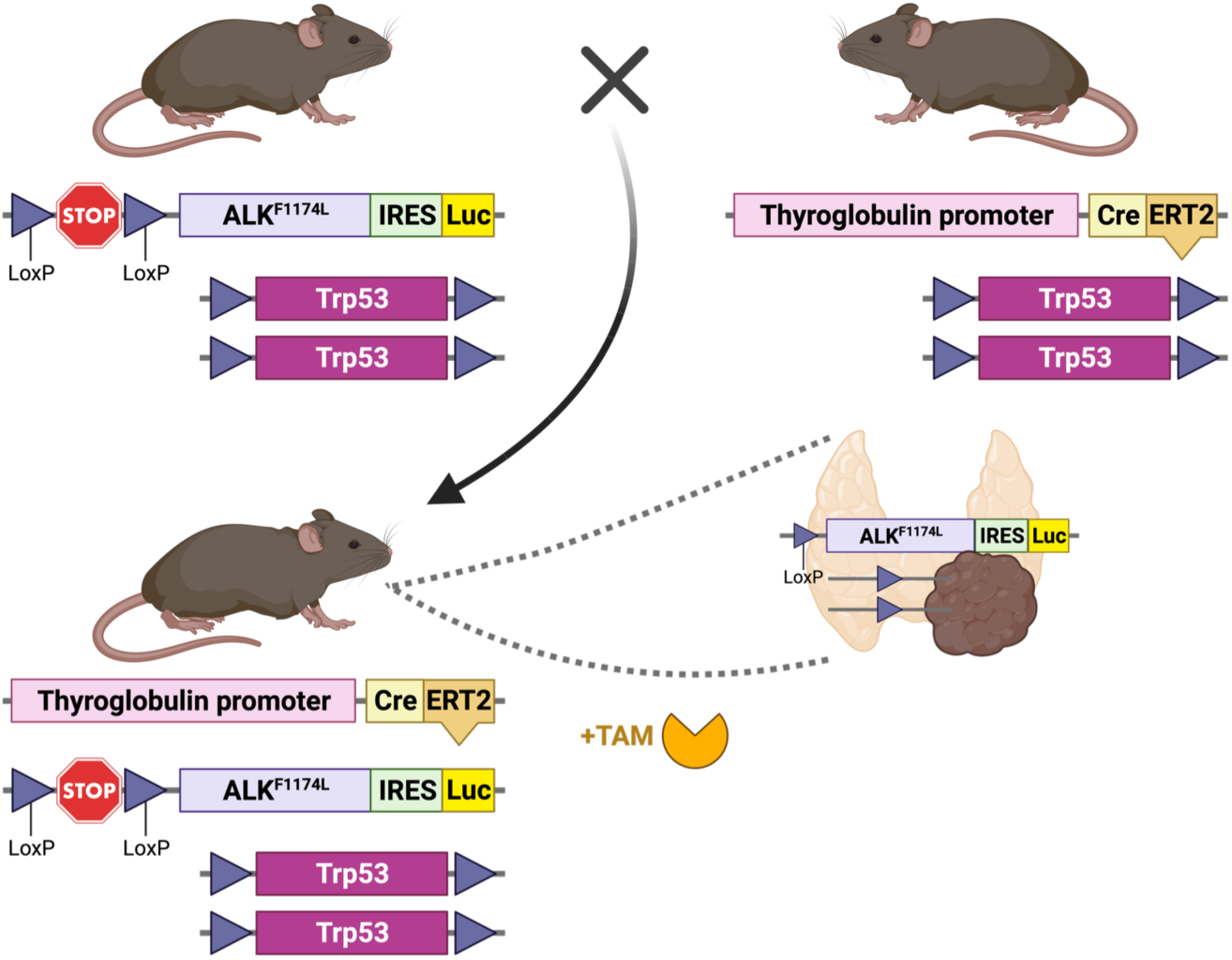
Generation of Trp53^KO^/ALK^F1174L^ mice. *Trp53*-floxed mice carrying the ALK^F1174L^ mutation were crossed with *Trp53*-floxed mice expressing CreERT2 recombinase under the thyroglobulin promoter to generate Trp53^KO^/ALK^F1174L^ mice. After tamoxifen (TAM) administration, the CreERT2 recombinase is specifically activated in thyroid follicular cells, leading to the expression of ALK^F1174L^ and the deletion of *Trp53*.

**Figure S2.**
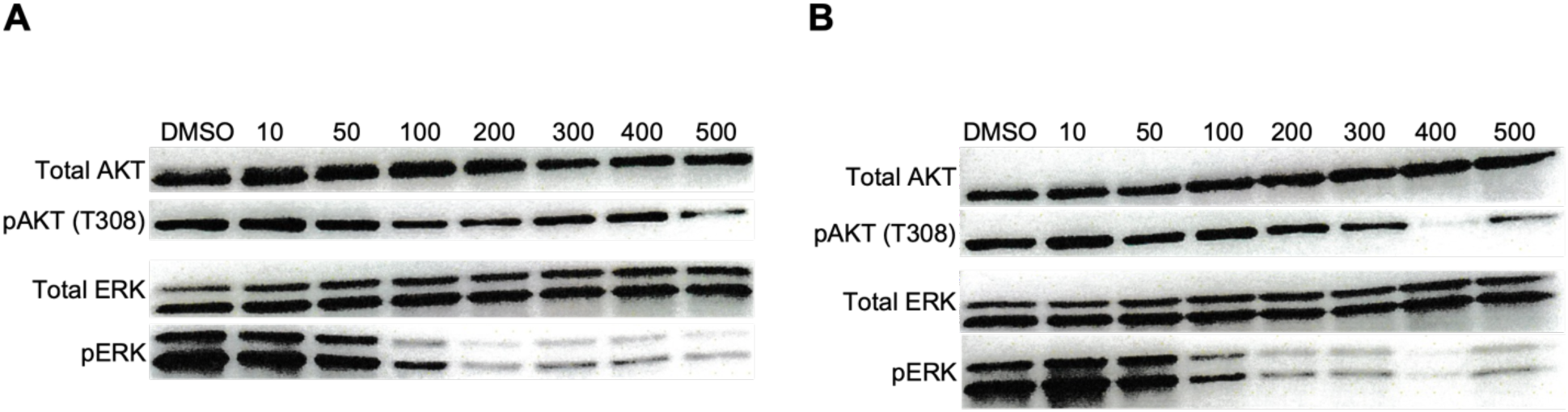
ALK inhibition in Trp53^KO^/ALK^F1174L^-derived PTC cell lines. Western blots showing that treatment with TAE-684 reduces phosphorylation of AKT and ERK in Trp53^KO^/ALK^F1174L^-derived PTC cell lines. Cells were derived from murine thyroids sacrificed on day 74 (**A**) and day 105 (**B**) after tamoxifen administration. Concentrations of TAE-684 are indicated in nM.

**Figure S3.**
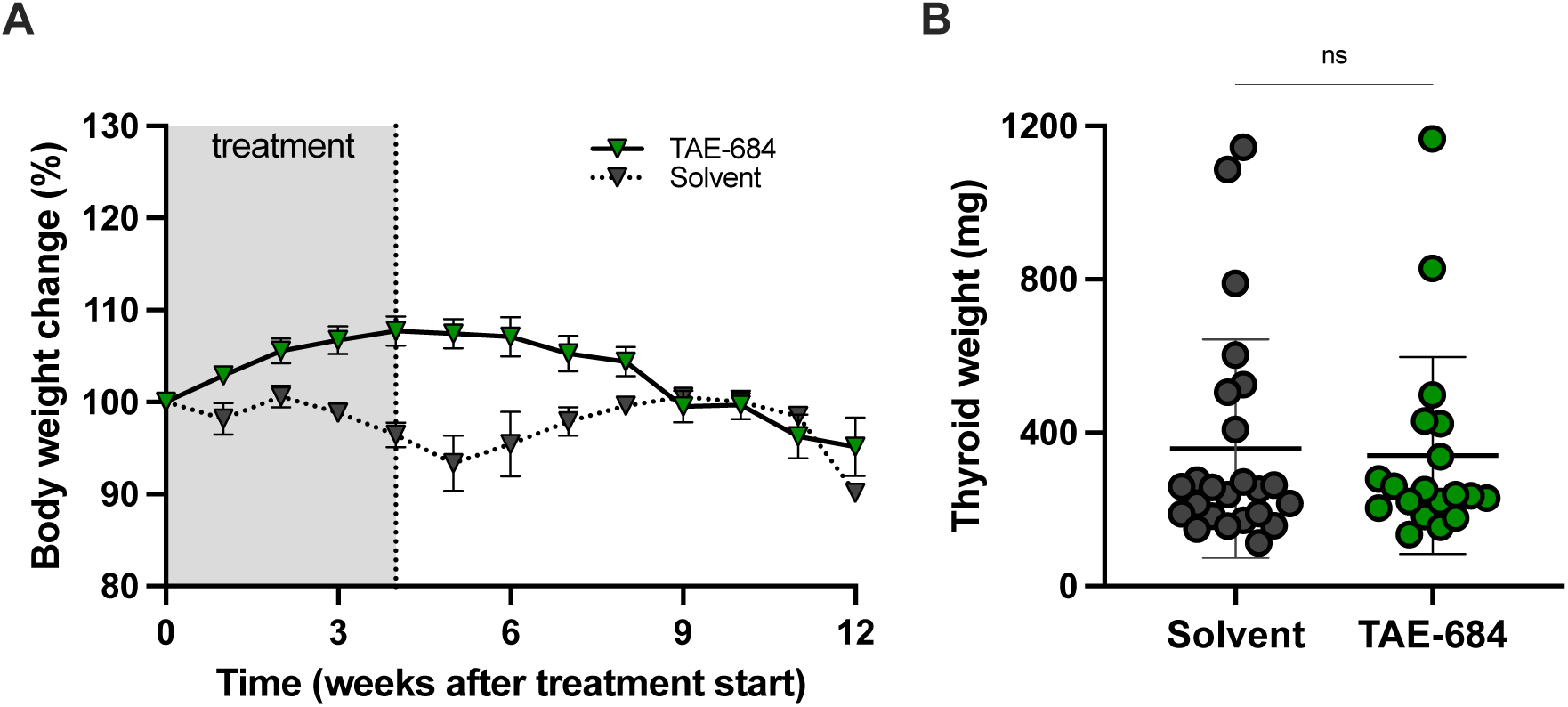
Effects of ALK inhibitor TAE-684 on body weight and thyroid tumour weight of Trp53^KO^/ALK^F1174L^ mice. **(A)** Changes in total body weight of TAE-684-treated Trp53^KO^/ALK^F1174L^ mice compared to the solvent-treated cohort (mean ± SEM). (**B**) Thyroid tumour weights at sacrifice show no difference between TAE-684-treated und solvent-treated mice. Ns, not significant.

**Table S1.** Score sheet for assessing the burden in Trp53^KO^/ALK^F1174L^ mice.

| **<u>Observation</u>** | **<u>Point assessment</u>** |
| --- | --- |
| **Body condition score (BCS)** |  |
| BC1: The animal is emaciated | 20 |
| BC2: The animal is underconditioned | 10 |
| BC3: The animal is in well-conditioned | 0 |
| BC4: The animal is overconditioned | 5 |
| BC5: The animal is obese | 10 |
| **Body weight** |  |
| 0-5% reduction or increase | 0 |
| Weight reduction 5-10% | 5 |
| Weight reduction 11-19% | 10 |
| Constant weight but with a sunken back | 10 |
| Weight reduction ≥ 20% | 20 |
| **General condition** |  |
| Shiny and clean coat, body openings clean, eyes clear and shiny | 0 |
| Mild constipation or diarrhea | 5 |
| Matted, dirty coat, sticky or wet body openings, abnormal posture, cloudy eyes, high muscle tone, mild dehydration (skin tenting resolves slowly) | 10 |
| Seizures, paralysis (trunk or limb muscles), apathy, breathing noises, cold to the touch, severe dehydration (skin tenting remains), severe constipation or diarrhea | 20 |
| **Spontaneous behaviour** |  |
| Normal behaviour (sleeping, reacting to cage movements and handling, curiosity, social interaction) | 0 |
| Slight deviation from normal behaviour | 1 |
| Unusual behaviour, restricted mobility, or mild hyperactivity | 5 |
| Self-isolation, lethargy, marked hyperactivity, or behavioral stereotypes, coordination issues, distended abdomen | 10 |
| Pain vocalizations when handled, self-amputation (automutilation) | 20 |
| **Clinical findings** |  |
| Breathing normal, limbs warm | 0 |
| Palpable enlargement of the thyroid gland | 10 |
| Noticeable increase in respiratory rate, tumour-related difficulty in consuming solid pellets, but softened food is still consumed | 10 |
| Gasping for air, tumour-related severe impairment of food intake such that even softened food is barely consumed | 20 |
| Tumour size $\geq 1000 \text{ mm}^3$, ulcerated tumour | 20 |

